# Sex-specific migration strategies of ospreys (*Pandion haliaetus*) from Germany

**DOI:** 10.1101/398735

**Authors:** Bernd-Ulrich Meyburg, Dietrich Roepke, Christiane Meyburg, Rien E. van Wijk

**Affiliations:** BirdLife Germany (NABU), PO Box 330451, Berlin 14199, Germany, ORCID 0000-0003-1673-4650; Gerhart-Hauptmann-Allee 26, 17192 Waren (Müritz), Germany; World Working Group on Birds of Prey, 31, Avenue du Maine, Paris 75015, France; Van Wijk Eco Research, Magle Torv 4, Søborg, Denmark, ORCID 0000-0002-6887-3153

**Keywords:** Bird migration, Sex-specific strategies, osprey, GPS tracking

## Abstract

Birds exhibit a wide variety of migration strategies, not only between species, but also within species. Populations might migrate to specific sites outside of the breeding season, but individuals within populations may also exhibit different migration strategies. Young, unexperienced birds may take different routes, visit different sites, and time their annual cycle differently than adults. In turn, within groups of adult birds, there may be a division between the sexes whereby males and females migrate to different sites or, more commonly, at different times. We investigated differences in the migration strategies of male and female ospreys (*Pandion haliaetus*) from a breeding population in northeastern Germany. An important difference between the sexes was the much earlier leaving of the breeding place by the females compared to the males. The difference in the timing of departure was much more pronounced compared to most other raptor species. The other main difference between the male and female ospreys was the distance that the birds accumulated over the annual cycle, with males generally moving more while at the breeding ground compared with the non-breeding grounds, and the opposite in females. An exception to this observation was two males that migrated to the Iberian Peninsula, that covered longer distances during the non-breeding season. Consequently, individuals accumulated the same distances over the course of an annual cycle, regardless of sex or migration strategy. Unexpectedly, a difference in the timing of the annual cycle between the sexes occurred at the breeding grounds with females leaving 2 to 3 months before the males, long before the young had fledged. Because males migrated much faster and, unlike the females, did not make prolonged stops, their arrival times at the non-breeding grounds were not different. Return migration to the breeding grounds was very similar between the sexes, and even the two males that spent the non-breeding season on the Iberian Peninsula did not arrive before any of the other birds. Thus, a shorter migration distance is not necessarily associated with an advantage with respect to a timely return to the breeding grounds.

## Introduction

Migration strategies of birds vary widely between species from short-distance, altitudinal migrations [1] to cross-hemisphere migrations of birds weighing only 25 g between breeding grounds in the northern hemisphere and non-breeding grounds in sub-Saharan Africa [2]. Not only are there differences between species, but even within species there might be population-specific migration routes [3] and non-breeding sites [4]. Even populations of the same species may spend the non-breeding season on different continents [5, 6]. Such differences within populations can relate to several factors, but are most commonly attributed to age and/or sex differences [7]. Juvenile, unexperienced birds are more sensitive to adverse weather conditions [8], have a lower navigation capacity [9], and often spend more time on stopovers compared with adults [10], making their annual cycles longer in both space and time than that of experienced adult birds. In adults, migration strategies often differ between males and females, especially in bird species that are also territorial at the non-breeding grounds or where there is pressure to return to the breeding grounds in a timely manner to secure the best territories and ensure high breeding success. For example, compared with Savannah sparrow (*Passerculus sandwichensis*) females, the males spend the non-breeding season closer to the breeding grounds, presumably because they are the dominant sex and strive to arrive earlier at the breeding grounds by minimizing the distance traveled to the non-breeding grounds [11]. An earlier return of the males than the females to the breeding grounds is a rather common phenomenon across migrating bird species [7].

In raptors, unlike most other bird species, females are the larger sex [12, 13]. Accordingly, among migratory raptors having a large size sexual dimorphism, females rather than males may spend the non-breeding season closer to the breeding grounds. The osprey (*Pandion haliaetus*) is a species that has been studied intensively, mainly using ringing recoveries. Female ospreys are larger than males, but the size difference between the sexes is relatively small [14], and thus differences in their migration strategies may also be minimal. In addition, ospreys are usually monogamous, occupying the same nesting site each year, and therefore the pressure to arrive early might not be as strong as for other species that must strongly compete for both females and nesting sites. Nevertheless, the ringing data of ospreys suggests that males do arrive earlier at the breeding grounds, partly because they spend the non-breeding season further north, closer to their breeding grounds [15, 16]. These findings, however, relate mainly to young birds and data on the migration strategies of adult ospreys from Central Europe is lacking. Here we investigated the migration strategies of a population of ospreys from northeastern Germany that was tracked between 1995 and 2017 using telemetry data with a special focus on potential differences between sexes.

## Materials and Methods

### Study site and data collection

We deployed satellite transmitters (PTTs) on 28 adult ospreys between 1995-2011 in northeastern Germany (25 in Mecklenburg-Vorpommern, 3 in Brandenburg) using a harness fixing the transmitter near the center of lift to minimize effects on the bird’s behavior [17, 18]. Ospreys were trapped by the so-called Dho Gaza method, most successfully using live adult sea eagles (*Haliaeetus albicilla*). The trapping, deployment of the transmitters and the ringing of the ospreys were approved by the competent authority in Güstrow (Landesamt für Umwelt, Naturschutz und Geologie, Agency for Environment, Nature Conservation, and Geology) - the exact name has changed a few times over the years - and the marking with the transmitters was treated in the same way as the ringing. Both nature conservation and animal welfare aspects were taken into account in the approval procedure, so that there was no separate procedure and no separate approval from the point of view of animal welfare.

### Transmitters

In 1995, seven battery powered PTTs from Microwave Telemetry Inc. (Columbia, MD, USA) were used. The lifetime of these transmitters is a maximum of 1 year when programmed for only a few hours at intervals of several days. In the following years up to 2005, 35g solar-powered PTTs (Microwave Telemetry, Inc.) that could be located by ARGOS using the Doppler phenomenon were deployed. The addition of a high-efficiency solar array freed the transmitter from the lifetime limitations imposed by a primary lithium battery. The lifespan of these transmitters is several years and with sufficient charge the batteries could record thousands of locations in total, but the fixes remained too inaccurate and irregular for the analysis of small-scale movements.

Since 2006, 30g solar ARGOS transmitters with GPS positioning that also transmit data on flight altitudes, direction, and speed, have been used. A total of 17 individuals were equipped with these tags. The number of fixes for these tags largely depended on how well the batteries were charged. Because ospreys usually sit on treetops or masts and flee a lot, the transmitters were usually sufficiently charged and recorded positions with an hourly interval between 4 am and 9 pm local time.

The complete mass of each of the transmitters and harnesses used in the present study was less than 2.2% of the body mass of the trapped bird and therefore only slightly impaired the bird’s performance. For further details, see Meyburg and Fuller (2007) and Meyburg and Meyburg (2013).

### Migration data

We obtained data of 8 individuals with ARGOS data totaling 13 bird-years and GPS data of 17 individuals totaling 33 bird-years (one individual being tracked for 8 consecutive years). In this paper, we focus only on the first year a bird was tracked. For fine-scale analyses on migration progression, we focused on the more axccurate and frequently recorded GPS dxata of the transmitters used since 2006.

### Data analysis

We cleaned up movement data for each individual, using the general purpose filters available in the R package *move* whereby we removed duplicate entries for the same individual with the same timestamp and positions with an average speed higher than 35 m^-s^. From the ARGOS satellite transmitter data, we additionally excluded positions with the location class “Z” (invalid locations). After filtering the data, we divided it into stationary versus movement periods. We defined a stationary period as a cluster of locations within a 30-km radius for at least 72 h. Using a moving window, points were added to the cluster until at least two consecutive points fell outside of the cluster. Next, clusters of locations were merged if the individual returned to a previous cluster for at least 72 h (e.g., if an individual returned to the breeding grounds after a long foraging trip). We could then define the timing schedule for each individual with a departure date from the breeding grounds, possible stops during migration, arrival date at the non-breeding grounds, departure from the non-breeding grounds and, finally, return to the breeding grounds.

To investigate diurnal and annual movements, we investigated several aspects: 1) the accumulated distances ospreys traveled during the breeding and non-breeding season, 2) the accumulated distances ospreys traveled over the annual cycle, 3) the mean distances of hourly movements, and 4) mean distances per hour over the course of the day for both the breeding and non-breeding season. To investigate aspects 1 and 2, we only used ospreys with data for a full breeding or non-breeding season or a complete annual cycle, whereas for aspects 3 and 4 we also used individuals with incomplete data for the entire annual cycle. To calculate the distances between consecutive locations, we used the package *geosphere* in R to make a distance matrix for each individual. For aspects 1 and 2, we accumulated the distances between consecutive positions for the respective periods and used simple linear models to test whether the accumulated distances differed between sexes and/or were a result of the number of locations, the number of observations per day, or the duration of the period. To investigate aspects 3 and 4, we followed the same approach to obtain distance matrixes, after which we only included positions between 4 am and 10 pm using the filter function in the *dplyr* package in R. We then selected only those consecutive positions with 1 h between them using the timeLag function in the *move* package in R. Finally, we excluded movements of more than 20 km between consecutive locations within either the breeding or non-breeding season (excluding <1% of all positions). To visualize aspect 3, we determined the density of movements in km^-h^ over classes of 1 km^-h^ (0-1 km^-h^, 1-2 km^-h^, etc.).

## Results

### Spatial migratory behavior

The ospreys migrated over a broad front southward to their non-breeding grounds, which were mainly located throughout western sub-Saharan Africa, but three males spent the non-breeding season on the Iberian Peninsula (Figure 1). Most of the birds spent the non-breeding season at the coast of Senegambia and Guinea-Bissau, but some migrated as far east as Lake Lagdo in northern Cameroon (Figure 1).

**Figure 1.**
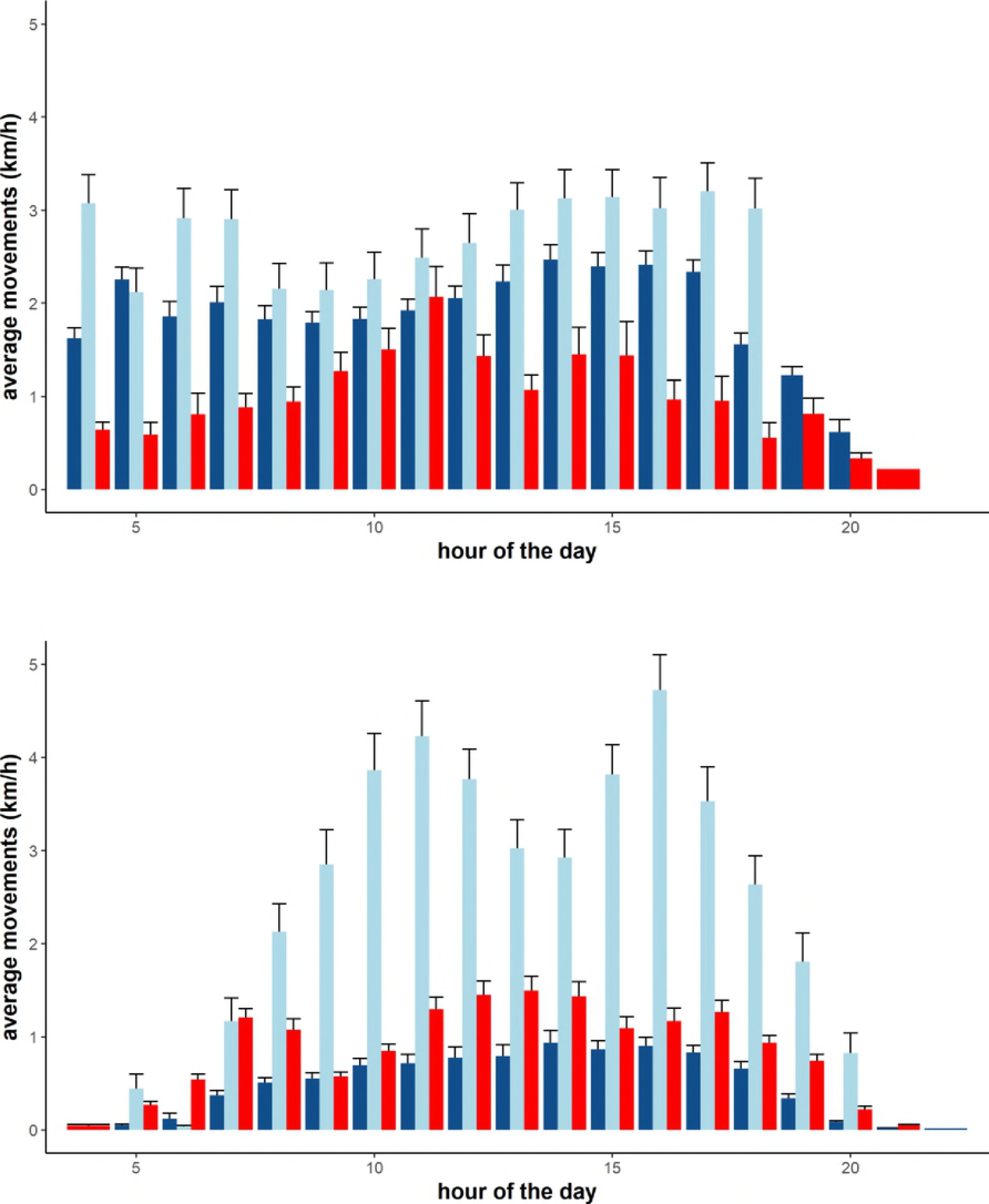
Autumn (left) and spring (right) migration routes of female (red) and male (blue) ospreys. Points indicate non-breeding sites. Individuals with only non-breeding sites had insufficient data on their exact migration routes.

### Annual and diurnal movements

Interestingly, males that migrated short-distances to the Iberian Peninsula accumulated the largest distances at both the breeding and non-breeding grounds (Figure 2). Females covered only short distances during the breeding season, but more so at the non-breeding grounds (and during migration), whereas long-distance males accumulated greater distances at the breeding grounds instead of the non-breeding grounds (Figure 2). Over the whole annual cycle, the short-distance males accumulated 13.897 to 17.315 km, long-distance males accumulated 16.295 km, and females accumulated 14.304 to 19.426 (on average 16.386) km.

**Figure 2.**
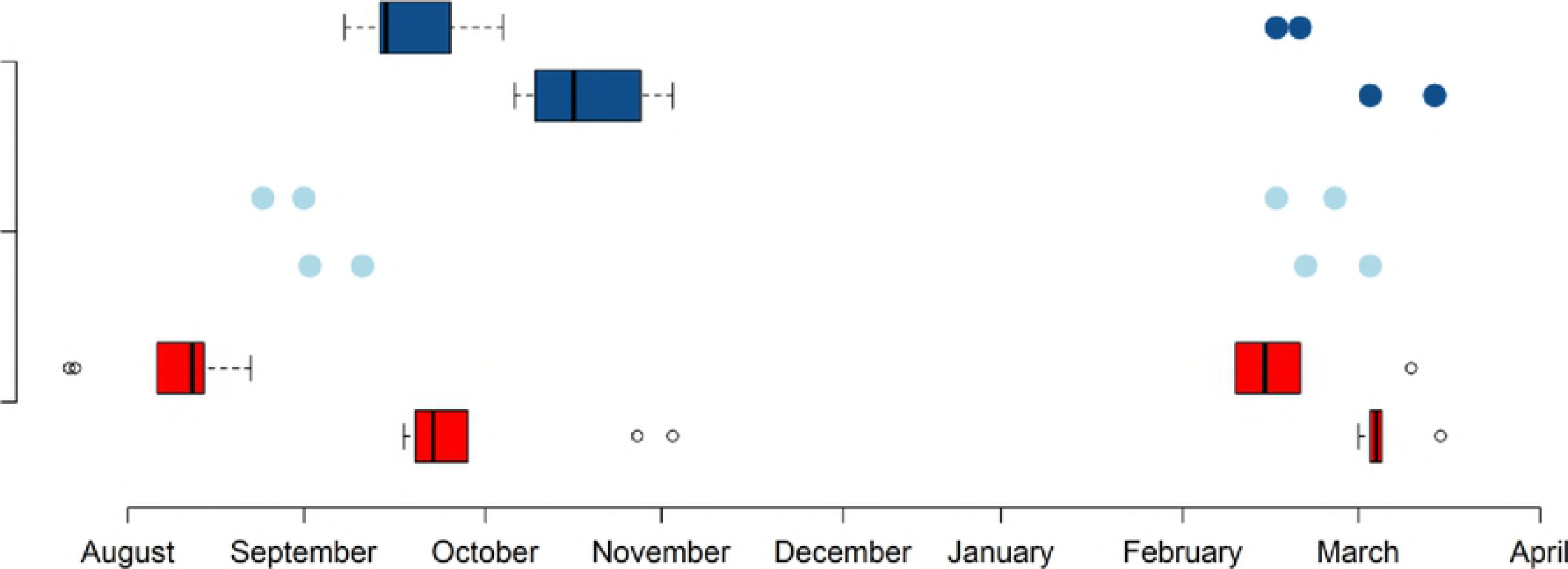
Accumulated movements within the breeding (left panel) and non-breeding season (right panel) for the different sex and migration classes. Dark blue represents males that migrated long-distances, light blue represents males that migrated short-distances, and red represents females.

These differences also translated to differences in the distances of hourly movements: for females, these were generally below 1 km during the breeding season, and for males, slightly longer (Figure 3). During the non-breeding season, there were no apparent differences (Figure 3), but there were differences in the daily patterns of movements: males migrating to sub-Saharan Africa did not exhibit distinct activity peaks, whereas the two birds that migrated to the Iberian Peninsula exhibited distinct activity peaks in the late morning and afternoon (Figure 4).

**Figure 3.**
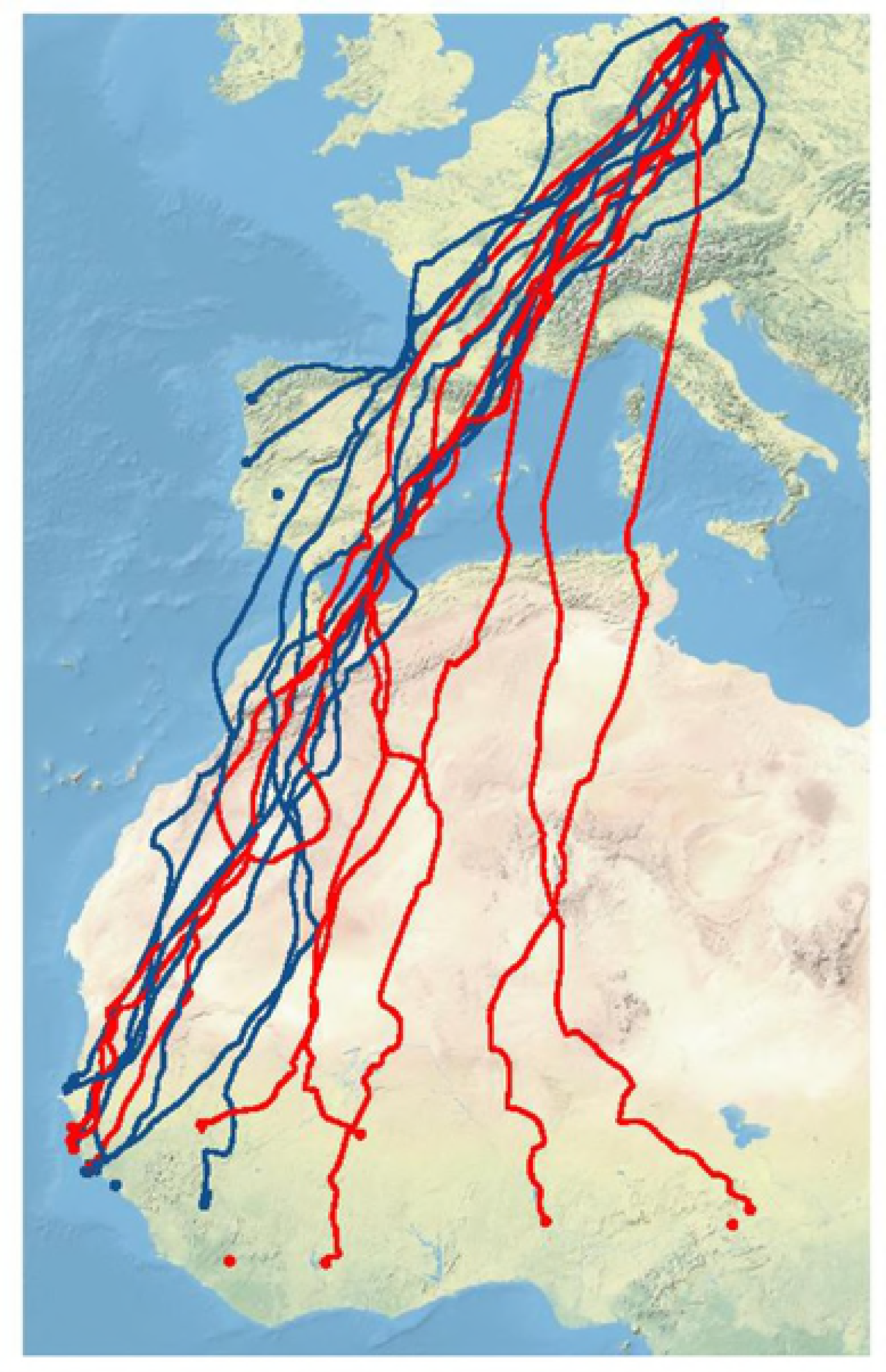
Density distribution of mean hourly distance during the breeding (upper panel) and the non-breeding season (lower panel). Females are represented in red, males that migrated long-distances to sub-Saharan Africa in dark blue, and males that migrated to the Iberian Peninsula in light blue.

**Figure 4.**
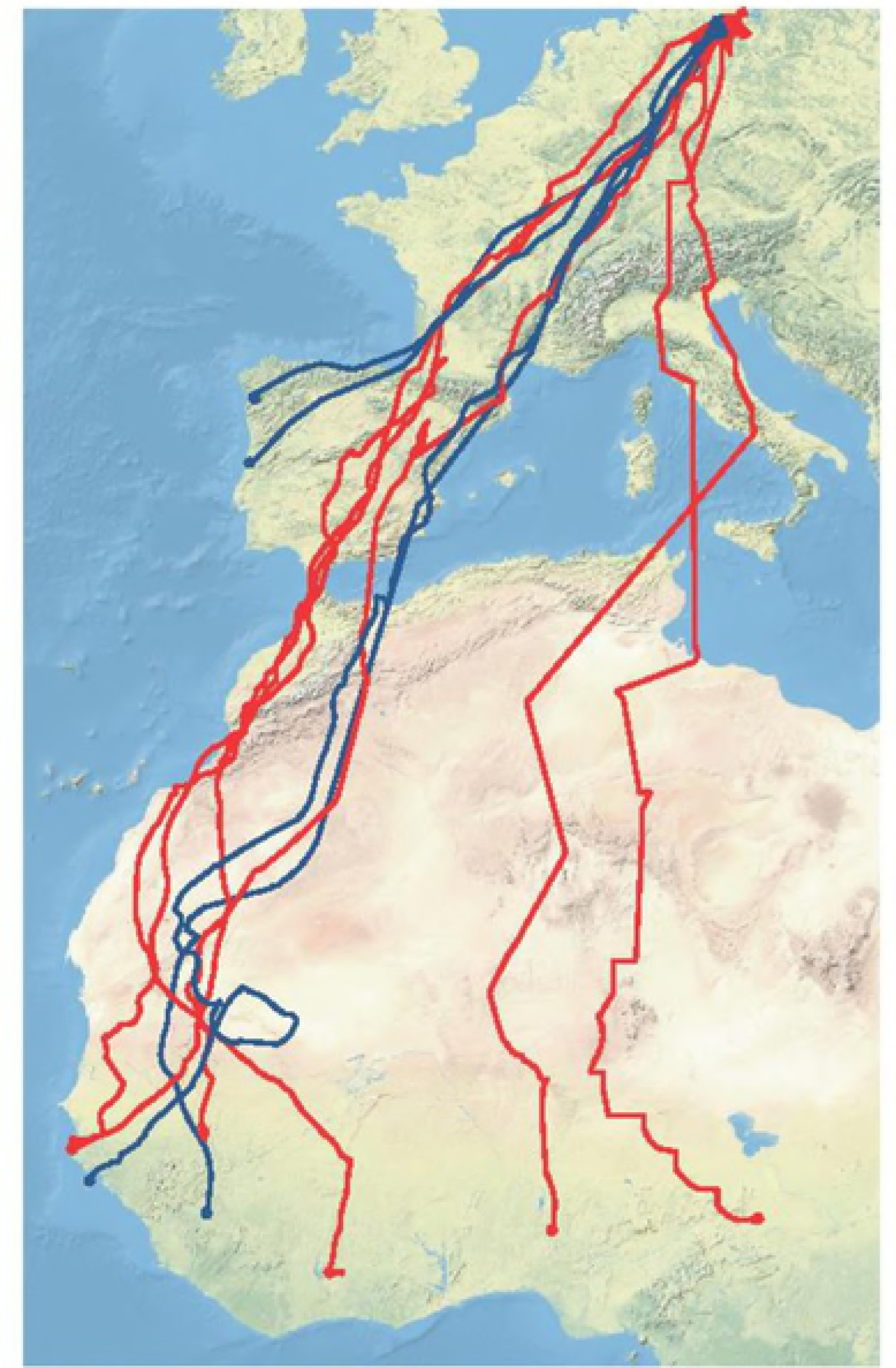
Distribution of the mean distance of movements per hour over the course of the day for the breeding season (upper panel) and the non-breeding season (lower panel). Females are represented in red, males that migrated long-distances to sub-Saharan Africa in dark blue, and males that migrated to the Iberian Peninsula in light blue. Means with standard errors are shown.

### Timing of migration

Female ospreys showed a form of ‘reversed protandry’, leaving the breeding grounds before the males (Figure 5). The males stayed at the breeding grounds 2 to 3 months longer until after the young had fledged but migrated faster and therefore ended up at the non-breeding grounds at around the same time as the females (Figure 5). Even though the two males that migrated to the Iberian Peninsula had to cover much shorter distances and arrived the earliest at their non-breeding grounds, they did not arrive earlier at the breeding grounds (Figure 5).

**Figure 5.**
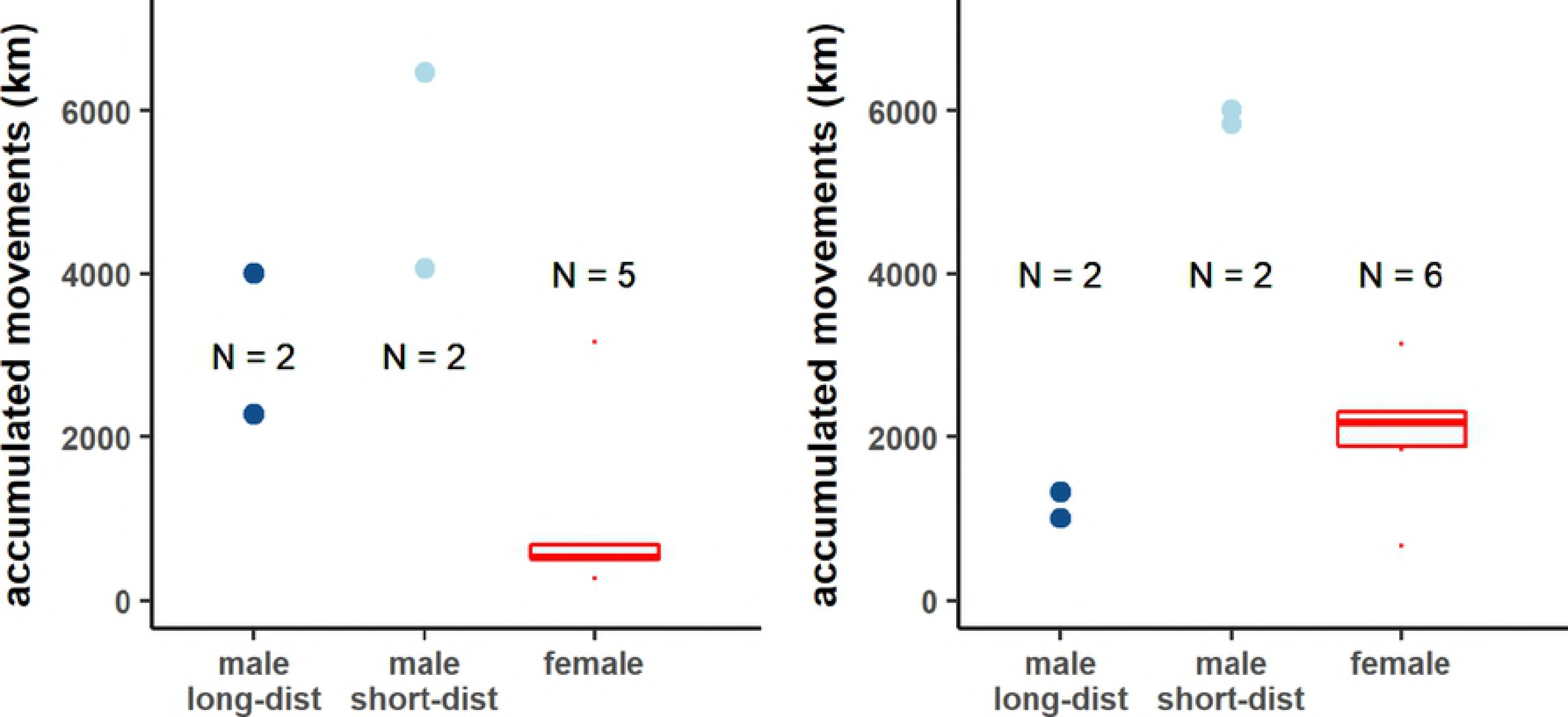
Timing of the annual cycle for males that migrate to sub-Saharan Africa (dark blue), males that migrate to the Iberian Peninsula (light blue), and females (red). Date of departure from the breeding grounds, arrival in the non-breeding grounds, departure from the non-breeding grounds, and arrival back in the breeding grounds are shown.

## Discussion

Ospreys from the studied population did not exhibit strong differences between the sexes regarding where they spent the non-breeding season or when they returned to the breeding grounds, but there were large differences in the timing of departure from the breeding grounds with females leaving months before the males. The other marked differences were in the distances accumulated over the different seasons with long-distance males covering considerably longer distances during the breeding season compared with the non-breeding season. Two males that migrated short-distances to the Iberian Peninsula did not show such a contrast.

Female ospreys are believed to stay for a long time near the nest after the hatchlings fledge [19]. Our results however show the opposite. Females in our study seem to leave the parental care fully to the males before the hatchlings have fledged; females leave the breeding grounds well before this time. Similar observations have been made before in other osprey populations [20], but not with such extreme differences. Interestingly, arrival times at the non-breeding grounds did not differ between the sexes since the females made prolonged stopovers of several weeks just south of the breeding grounds. Males most likely manage to build up body reserves for migration while feeding the young.

How families time their departure, has been little studied to date in larger raptors. In Greater Spotted eagles (*Clanga clanga*), it was shown that the female left the breeding site only two to three days before the fledglings, the male left one week later [21]. The reasons for this contrasting behavior between species, are hardly known. Since ospreys kill relatively large prey, it should not be a problem for a male to feed even up to three young alone, which is in contrary to e.g. a bigger eagle such as the Greater Spotted eagle that must raise offspring with much smaller prey items. Female ospreys seem to take advantage of this by leaving earlier, leaving post-fledgling care solely to the male. Some females even left the eyrie for extensive foraging trips over 50km away already way before young had fledged, an observation that has been noted in females of the smaller Lesser Spotted eagle (*Clanga pomarine*) as well [22]. An additional explanation why females leave before males and juveniles, can relate to the fact that they can already start moulting feathers during incubation, giving them a head start, but future comparisons in the moulting schemes of ospreys with contrasting migratory behavior across their breeding range can hopefully shed more light on that [23].

These findings highlight differences in the timing of the annual cycle and differences in biology between males and females. Whereas females are the parent mostly responsible for incubating the eggs [24, 25], the male provides food throughout this time. Until the hatchlings are a few weeks old, only the male provides the food [26]. Furthermore, even though females engage in territorial defense [24], it is mainly the males that protect against possible intruders [27]. Therefore, the males naturally covered more distance at the breeding grounds. Likewise, because the females do not have to search for food during the breeding season but do have to search for food at the non-breeding grounds, they accumulate much longer distances during the non-breeding season. Males on the other hand, seem to reduce their movements at the non-breeding grounds, except for the two males that migrated short distances.

At the non-breeding grounds, the most marked difference was the reduction in the length of hourly movements, indicating that individuals minimize the distance between resting and foraging grounds. Males that migrated to the Iberian Peninsula exhibited a different pattern with two distinct activity peaks and longer movements. Personal observations confirm this pattern with the males in this area moving out from an inland forest to the coast to forage both in the morning and evening (Meyburg et al. *pers. obs.*). Presumably, these distances are covered to avoid kleptoparasitism by local gull species (*Larus sp.*). This may indicate that food is not as readily available in this area compared with the sub-Saharan non-breeding grounds. Consequently, migrating short-distances might at first seem advantageous, but in the long run, short distances require similar or even greater energy investments than long-distance migrations. Similar to findings in Lesser black-backed gulls (*Larus fuscus*, [28]), in end effect, the total accumulated distance over the year, is the same regardless of the individual’s sex or migration strategy.

## Declarations

### Acknowledgements

The permits for the capture of the birds and their fitting with transmitters were issued by the state ministries of the federal states of Mecklenburg-Vorpommern and Brandenburg, Germany. The Hiddensee ringing station supplied data for trapped birds that were already ringed. For assistance and advice concerning the satellite transmitters, we wish to thank Dr. P. Howey (Microwave Telemetry Inc.) for his kind support. Michael McGrady kindly proofread the manuscript.

### Author contributions

Conceptualization of the study – B-UM, CM, DR, AB, RVW

Methodology – B-UM, CM, DR, AB, RVW

Statistical analysis and visualization – CM and RVW

Data analysis – B-UM and RVW

Resources – B-UM

Data curation – CM

Writing – B-UM and RVW

Project administration and funding acquisition – B-UM

### *Permit*s

The permits for the capture of the birds and their fitting with transmitters were issued by the state Ministries of the federal states of Mecklenburg-Vorpommern and Brandenburg, Germany.

